# The Role of Low Molecular Weight Fungal Metabolites in Grapevine Trunk Disease Pathogenesis: Eutypa Dieback and Esca

**DOI:** 10.1101/2021.08.09.455770

**Authors:** Gabriel Perez-Gonzalez, Dana Sebestyen, Elsa Petit, Jody Jellison, Laura Mugnai, Eric Gelhaye, Norman Lee, Sibylle Farine, Christophe Bertsch, Barry Goodell

## Abstract

Eutypa dieback and Esca are serious grapevine trunk diseases (GTDs) caused by fungal consortia causing large economic losses in vineyards. Depending on the disease the species involved include *Eutypa lata, Phaeoacremonium minimum*, and *Phaeomoniella chlamydospora*. There is a need to understand the complex pathogenesis mechanisms used by these causative fungi to develop treatments for the diseases they cause. Low molecular weight metabolites (LMW) are known to be involved in non-enzymatic oxygen radical generation in fungal degradation of wood by some Basidiomycota species, and as part of our work to explore the basis for fungal consortia pathogenesis, LMW metabolite involvement by the causal GTD fungi was explored. The GTD fungal pathogens examined, *Eutypa lata, Phaeoacremonium minimum* and *Phaeomoniella chlamydospora*, were found to produce low molecular weight iron binding metabolites that preferentially reduced iron or redox cycled to produce hydrogen peroxide. Uniquely, different LMW metabolites isolated from the GTD fungi promoted distinct chemistries that are important in a type of non-enzymatic catalysis known as chelator-mediated Fenton (CMF) reactions. CMF chemistry promoted by LMW metabolites from these fungi allowed for the generation of highly reactive hydroxyl radicals under conditions promoted by the fungi. We hypothesize that this new reported mechanism may help to explain the necrosis of woody grapevine tissue as a causal mechanism important in pathogenesis in these two grapevine trunk diseases.

**IMPORTANCE:** Understanding the pathogenesis of grape trunk diseases (GTDs) is the key to the development of disease control and treatment. While fungal extracellular enzyme systems are typically cited relative to their fungal mechanisms in pathogenesis, non-enzymatic mechanisms have been less studied in this regard and the role of low molecular weight (LMW) fungal metabolites in GTD development is quite limited. In this article, we demonstrate that GTD-causative fungal pathogens *Eutypa lata, Phaeoacremonium minimum* and *Phaeomoniella chlamydospore* produce LMW phenolic metabolites under iron-restricted conditions. These metabolites undergo a series of redox reactions, with different fungi producing metabolites that preferentially either reduce iron, or generate hydrogen peroxide, under conditions simulating grapevine woody tissue. These conditions have the potential to promote generation of highly damaging hydroxyl radicals through a mechanism that appears to be similar to non-enzymatic chelator-mediated Fenton (CMF) chemistry which is involved in fungal degradation of wood by non-related fungal orders. This is the first report of CMF chemistry promoted by GTD-causative fungi under laboratory conditions and the research suggests an alternate pathway that may contribute to pathogenesis in GTDs, and a potential target for vine protection.

## INTRODUCTION

Grapevine trunk diseases (GTDs) are caused by a complex of fungi that were described as early as the end of the 20th century. Most GTD fungi attack the perennial tissues of the grapevine and ultimately lead to the death of the plant (1-3) and are characterized by the dieback and necrosis/decay of the stem tissue. Some of these diseases can show foliar symptoms that may not appear until deterioration of the stem wood is advanced (4). Although Eutypa dieback has been reported to be caused solely by *Eutypa lata* or other *Eutypa* species, current literature suggests that these pathogens are also associated with other GTDs. Further, *Eutypa* spp. is often associated with a consortium of other fungi, especially the Diaporthales order, with *Phaeoacremonium minimum* (Pmin) and *Phaeomoniella chlamydospora* (Pch) predominating (5-8). Eutypa dieback, Esca, and *Botryosphaeria* dieback are the most significant GTDs involving one or several xylem-inhabiting fungi, with Pch and Pmin typically found in consortia for Esca disease development (3, 9).

Most GTD fungal pathogens enter grapevine trunk wood in vineyards through pruning wounds, inhabiting the xylem cells in the woody tissue and causing, with time, significant necrosis and decay, ultimately leading in some of those diseases to foliar symptoms and cordon and vine death (4). In Eutypa dieback, as the disease progresses complete loss of yield, stunting of shoots, and/or loss of cordons and vines occurs, with older vineyards experiencing as much as 30% necrosis of cordons or vines. In the USA, Eutypa dieback and *Botryosphaeria* dieback predominate in California and have also been an emerging issue for cold-climate vineyards in the Northeastern US and in British Colombia, Canada. In California alone, losses by GTDs each year amount to 14% of the value of the wine grapes produced with economic loss of more than $260 million per year (10, 11). GTDs also cause up to 2 billion in losses to vineyards globally each year (12).

While the pathogenic mechanisms and foliar damage in Eutypa dieback has been reported to be associated with the production of eutypine and other phytotoxic compounds (13), the mechanisms involved in producing woody tissue necrosis and other symptoms associated with both Eutypa dieback and Esca are still not well understood. Importantly, it is also unknown why a consortium of fungi is typically involved in both diseases (13, 14).

Wood decay by brown rot Basidiomycota species is similar in some respects to the necrosis in grapevine wood caused by Ascomycota GTD fungi. Decay initiated by brown rot fungi produces a type of wood degradation where both holocellulose and lignin are depolymerized by a highly oxidative non-enzymatic chelator mediated Fenton (CMF) mechanism (15-21). After, or concurrent with CMF action, the polysaccharide component of the wood cells is then preferentially extracted from the wood via the action of fungal extracellular carbohydrate active enzymes (CAZymes), while the lignin component is repolymerized within the wood cell wall (19, 22). In *E. lata*-infected grape wood, structural glucose and xylose from the hemicellulose are also preferentially degraded and depleted from the wood while lignin is degraded much more slowly (23) similarly to brown rot wood decay. The residual lignin in both brown rotted wood and in necrotic wood attacked by GTD fungi is brown in coloration because of the residual or modified lignin which remains. Because of the similarities to brown-rot wood decay, and prior reports on the action of oxygen radical generation in Esca disease (24), we considered that a mechanism similar to CMF chemistry might potentially play a role in GTD fungal attack of grapevine wood. The hallmark of the CMF system is the production of fungal low molecular weight (LMW) compounds that promote the mediated Fenton reaction (eq. 1) by the reduction of Fe^3+^ to Fe^2+^ with the wood cell wall, and away from the fungal hyphae (15, 19). This CMF chemistry is promoted by a reduction in pH of the fungal micro-environment (typically to pH 5.5 or lower). Brown rot fungi generally reduce the pH of their micro-environments to approximately pH 4 or lower, and this is thought to aid in promoting a sequence of reactions leading up to the Fenton reaction occurring within the wood cell wall (19, 25, 26). In the low pH environment of the brown rot fungal extracellular matrix (ECM) which immediately surrounds the fungal hyphae, the pH is lower than pH 4.0 and iron will be sequestered (19). However, within the wood cell wall the pH is maintained at approximately 5.5, and at that pH LMW compounds will redox cycle with iron to generate Fe^2+^ and also generate H_2_O_2_ (27, 28). Both of these reactants are required for the generation of hydroxyl radicals (HO^•^), which are required to be generated within the cell wall for oxidative depolymerization of both cellulose and lignin to occur. This action then leads to further cell wall deconstruction by fungal CAZymes (22, 29).

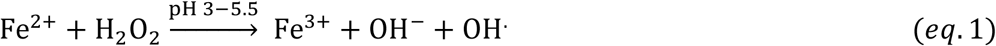

Osti and DiMarco reported that fungal supernatants of Pch and Pmin contained phenolate siderophore-like compounds, and their analysis suggested that catecholate compounds from the fungal supernatants were present that could also reduce iron, generate hydroxyl radicals (HO^•^) and depolymerize cellulose (24). However, their supernatants also contained high molecular weight components such as extracellular enzymes which could have skewed some of their results. Therefore, in our research we assessed only the ultrafiltered <5kDa fraction of LMW metabolites. We did not assess for siderophore receptor sites on the fungal membranes as examination of classic siderophore function was not part of our objectives for this research.

*Eutypa lata* (Elata) and other fungal species involved in GTDs are known to produce acetylenic phenols and heterocyclic analogues similar to some types of siderophores (30, 31), and several LMW phenolic metabolites have been identified from Pmin and Pch (32). But it has not been reported whether these compounds have the ability to redox cycle or reduce iron as has been observed with some types of catecholate compounds produced by brown rot decay fungi, such as 2,5-dimethoxyhydroquinone (2,5-dimethoxybenzoquinone) (15).

Our research examines the nature of LMW compounds produced by Elata, Pmin and Pch, and by combinations of these three fungi, specifically for iron reduction and hydroxyl radical generation to determine if there is a link between the LMW metabolites produced and non-enzymatic chemistries that may produce hydroxyl radicals in a manner similar to that which has been demonstrated in CMF chemistry with other wood degrading fungi. We hypothesize that chemistries similar to the CMF mechanism, and enhanced by LMW metabolites produced by these fungi, could potentially play a key role in the initial stages of Eutypa dieback and Esca disease, particularly associated with wood grapevine tissue necrosis.

## MATERIALS AND METHODS

The three fungi involved in the Eutypa dieback consortia used in our current work were *Phaeomoniella chlamydospora* (Pch, isolate UCD7872), *Phaeoacremonium minimum* (Pmin, isolate UCD7770), and *Eutypa lata* (Elata, isolate UCD7746); all isolated from vineyards in Lodi, California. These fungi were grown both in single culture and in combination (Elata_Pch; Elata_Pmin; Pmin_Pch) for six-weeks in low-iron media. LMW iron-binding metabolite production was assessed, as was hydroxyl radical generation. All analysis were done in triplicate unless otherwise stated.

### Culture media

Iron-free cultures with restricted nutrient media were used to promote biosynthesis of iron-binding LMW compounds (33), with all glassware acid washed in 10% HCl for 24 hours, rinsed with deionized distilled water (ddH_2_O - 18.2MΩ.cm) followed by a 90mM EDTA wash, and then rinsed 3 times with ddH_2_O. Restricted nutrient, iron-free media was prepared modified from (34) as follows: 1L of ddH_2_O mixed with 2g ammonium nitrate (Sigma-Aldrich, MO, USA), 2g monobasic potassium phosphate (Merck, MA, USA), 0.5g magnesium sulphate heptahydrate (Sigma-Aldrich, MO, USA), 0.1g calcium chloride (Bio Basics Canada Inc, ON, Canada), 0.57mg boric acid (Sigma-Aldrich, MO, USA), 0.31mg zinc sulphate heptahydrate (HIMEDIA labs, PA, USA), 0.039mg copper sulphate pentahydrate (Acros Organic, Belgium), 0.036mg manganese chloride tetrahydrate (Fisher Chemical), 0.018mg ammonium molybdate tetrahydrate (Acros Organic, Belgium), and 0.001g thiamine HCl (Acros Organic, Belgium). For carbon sources, glucose (Alpha Biosciences) 0.5% (w/v) and 50µm microcrystalline cellulose (Acros Organics, Belgium) 1% (w/v) were used with five replicates for each culture and in each medium. The media solution (200ml per 0.5L flask) was brought to a pH of 5.5 using NaOH and autoclaved for 30 minutes. Liquid cultures were inoculated using mycelial slurries prepared by scraping mycelium from fully grown agar plates into 50mL of sterile ddH_2_O. One mL of mycelial slurry was carefully pipetted onto the surface of the liquid media in each flask and allowed to grow for six weeks to promote production of LMW metabolites.

### LMW metabolite extraction

After incubation, inoculated liquid cultures were coarsely filtered with Whatman #4 cellulose filters to remove the mycelium and cellulose microcrystals from the liquid cultures, and this filtrate was then serially filtered through a 0.22μm cellulose filter under vacuum. This was then followed by ultrafiltration through a 5kDa polyethersulfone filter used with an Amicon Stirred Cell filtration unit (EMD Millipore, MA, USA) to yield the <5kDa LMW metabolite fraction. A Bradford assay was conducted to confirm that proteins were not present (results not shown). The LMW metabolite fraction was acidified to pH 3 with HCl prior to a triple ethyl acetate (1:1 with ultrafiltered media) extraction for phenolics (33). The organic fraction was dried under reduced pressure, resuspended in methanol, and filtered through a 0.22μm filter to yield the final <5kDa LMW extract.

### Determination of total phenols by the Folin-Ciocalteu assay

Folin-Ciocalteu (F-C) reagent was used to spectrophotometrically quantify (765nm) the amount of total phenols in solution (35) with gallic acid used as the standard. Samples of purified extract (20µl) and the F-C reagent (100µl) were reacted (5min @RT) before the addition of 20% sodium carbonate (300µl) to initiate the F-C reaction. After a further reaction period (2h @RT) in the dark, absorbance values were compared to the gallic acid standard for phenolic concentration determination.

### Determination of iron reduction by Ferrozine assay

The Ferrozine assay is a colorimetric assay that is sensitive specifically to the ferrous form of iron and it is therefore useful in measuring iron reduction capacity (36). Each reaction cuvette contained at their final concentration: acetate buffer (0.100mM, pH 5.5), Ferrozine (0.250mM), Fe^3+^ (0.030mM). Phenolics in the fungal extract (15mM according to F-C assay) was added last to start the reaction. Ferrous chloride (FeCl_2_) was used for the standard. Samples were mixed thoroughly into each cuvette and the reaction followed spectrophotometrically (562nm) at 5 min intervals for 45min. The ferric iron reductive capacity of all LMW extracts was normalized per nmol of phenolics added to the reaction and reported as the amount of iron reduced (mol)/amount of phenolic (mol) in the extracts assayed.

### FOX assay for H_2_O_2_ detection

A ferrous ammonium – xylenol orange (FOX) assay (37) was used to measure H_2_O_2_ evolution which occurs during the oxidation of ferrous iron in Fenton chemistry. The oxidized iron reacts with xylenol orange (XO) to form a blue purple complex that can be detected at 560nm (37, 38). MES buffer (50mM, 5.5pH) and phenolics in fungal extracts (15mM according to F-C assay) were added to 100μL of the FOX reagent (a 1:1 ratio of the XO/Ferrous ammonium sulfate mixture and sorbitol) to yield a solution of 0.100mM XO, 0.250mM ferrous ammonium sulfate, 25mM H_2_SO_4_, and 100mM sorbitol, which was incubated (RT, 30min) before centrifugation (10,000RPM, 5min) to remove any precipitate. (Sorbitol was added consistently to all samples to increase the yield of ferric iron). All samples were maintained in low light conditions to reduce UV interference, analyzed at 560nm for H_2_O_2_ detection and reported as the amount of H_2_O_2_ generated (mmol)/amount of phenolic (mol) in the extracts assayed.

### Electron paramagnetic resonance (EPR) for detection of hydroxyl radicals

Methanol was evaporated from each sample using a SpinVac (35°C, 45min) after adding the fungal extract into the reaction solvent at two pH values: (i) sodium acetate/acetic acid buffer (80mM, pH 5.5) and (ii) acidified water (adjusted to pH 3.5 with HNO_3_). For hydroxyl radical detection in EPR, the following components (final concentration) were used in the reaction mixture: DMPO spin-trap (5,5-dimethyl-1-pyrroline-n-oxide, 10mM) was added to the other reaction components: H_2_O_2_ (0.15mM), phenolics in the fungal extract (50mM adjusted per the F-C assay). Fe^3+^ (0.15mM) was added to start the reaction, and after incubation (5 min, @RT) the samples were transferred to a 50μL EPR capillary glass tube. Catechol (50mM) was used as reference compound. Analysis was conducted in the X-band frequency (9.8 GHz) using a Bruker Elexsys-500 EPR instrument equipped with a super high QE cavity (ER4122SHQE-W1).

### High performance liquid chromatography (HPLC)

LMW extracts were analyzed using a Shimadzu HPLC system with an analytical C18 Nucleosil column (250mm x 4.6mm x 5µm, 0.50mL/min). Flow-through UV analysis (280nm) was conducted using an acetonitrile (ACN) linear gradient (10-90%) over 45min. The gradient was then held at 90% for 5min and the gradient reversed over the next 5 min before ending at 10% ACN over a total 60min. Fractions were then scanned (190 – 370 nm) to detect potential phenolic compounds at ∼280nm absorption.

### Identification of metabolites by ultra-performance liquid chromatography – mass spectroscopy (UPLC-MS)

Fungal extracts were analyzed using an UPLC (ACQUITY HSS T3 (100×2.1mm×1.8μm) with Ultimate 3000 LC) combined with a Q Exactive MS (Thermo) and screened with electrospray ionization MS (ESI-MS). The mobile phase was: solvent A (0.05% formic acid water), and solvent B (acetonitrile) with a gradient elution (0-1.0min, 5%B; 1.0-12.0min, 5%-95%B; 12.0-13.5min, 95%B; 13.5-13.6min, 95%-5%B; 13.6-16.0min, 5%B). The flow rate of the mobile phase was 0.3mL·min^-1^ with the column temperature (40°C), and the sample manager (4°C), both constant.

Mass spectrometry parameters in ESI+ and ESI-mode were:

ESI+ : Heater Temp 300 °C; Sheath Gas Flow rate, 45arb; Aux Gas Flow Rate, 15arb; Sweep Gas Flow Rate, 1arb; spray voltage, 3.0KV; Capillary Temp, 350 °C; S-Lens RF Level, 30%.

ESI-: Heater Temp 300 °C, Sheath Gas Flow rate, 45arb; Aux Gas Flow Rate, 15arb; Sweep Gas Flow Rate, 1arb; spray voltage, 3.2KV; Capillary Temp,350 °C; S-Lens RF Level,60%.

Raw data was analyzed using Compound Discoverer™ (ThermoFisher Scientific) a high-resolution accurate-mass data software for metabolite identification, and a comprehensive review was conducted for each of the main compounds found in the extract selected by peak area or peak height.

## RESULTS AND DISCUSSION

Cultures where two fungi were grown in consortia in some instances maintained a cultural appearance that was characteristic of one of the species, and this always occurred when Pmin was part of the consortia. With the Elata_Pch consortia growth, discrete colonies of both species were apparent in about half the cultures whereas in the other half, Elata growth predominated. However, when comparing the metabolomic profile of the individual cultures to the consortia growth, unique extract profiles were observed in the consortia cultures as detailed below and in Fig. S1.

### Production of phenolics and iron reduction capacity

Preliminary chromatographic screening of all fungal extracts by HPLC showed that putative phenolic compounds absorbing in the 280 nm region were present (Fig. 1). All extracts contained a maximum of five detectable peaks absorbing in the 280 nm range. Fig. 1 shows only the two most prominent phenolic peaks for extracts from the three fungi.

**FIG 1.**
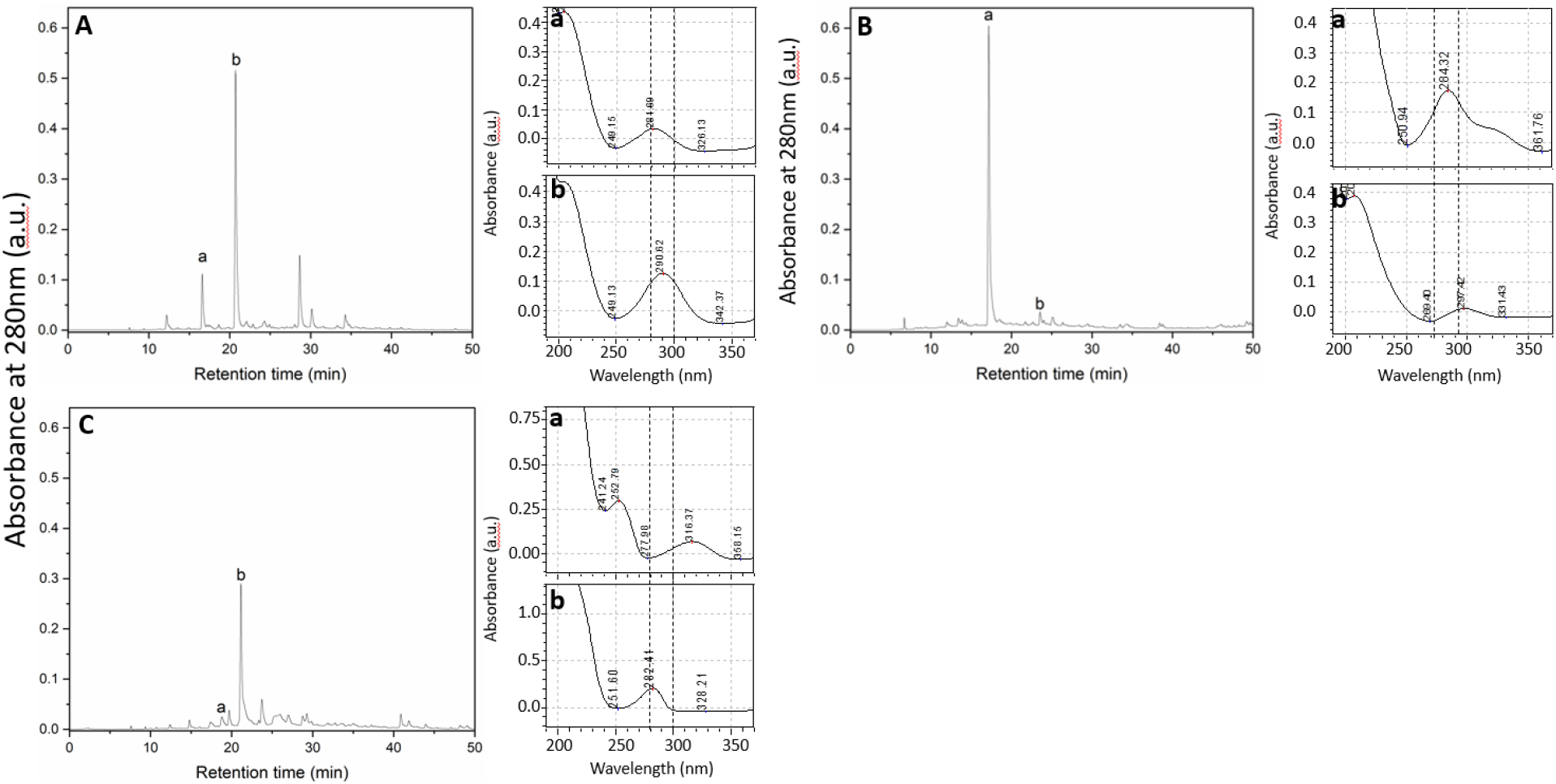
HPLC chromatograms (Left for each sample) of (A) Elata, (B) Pmin, and (C) Pch fungal metabolite extracts taken at 280nm. Labels (a,b) indicate the putative phenolic peaks selected (abs. ∼280nm). (Right) UV spectra of the two most abundant extract peaks.

The Folin-Ciocalteu (F-C) assay for phenolics in individual ∼280nm UV fractions showed that Pch extracts contained the largest amount of phenolics when compared to either the Elata or Pmin individual fungal culture extracts (Table 1, p < 0.001, p < 0.001). Extracts from all fungal consortia yielded an increase in total phenolics over individual cultures (Table 1). The Elata_Pmin consortia produced significantly more phenolics than either Elata or Pmin alone (Table 1, p = 0.01, p = 0.03). For the Elata_Pch consortia growth there was an averaging effect relative to phenolic production. Phenolic content when Elata and Pch were grown in combination was significantly lower than Pch alone (p < 0.001), but it was significantly higher than Elata alone (p = 0.007). The Pmin_Pch combination showed an increase in overall phenolic production compared to that when the fungi were grown individually. The additive effect alone may be an important finding relative to Eutypa dieback and Esca.

**TABLE 1.** Total phenolic content (Folin-Ciocalteu) of LMW fungal extracts from the three GTD fungi, together with iron reduction (Ferrozine), and H_2_O_2_ production (FOX) from those extracts. Results shown as mean ± SD, n = 3.

|  | *Total phenolic content*<br>(mM) | Iron reduced (mol<br>Fe<sup>2+</sup>/mol of phenols) | H<sub>2</sub>O<sub>2</sub> produced (mmol<br>H<sub>2</sub>O<sub>2</sub>/per mol phenols) |
| --- | --- | --- | --- |
| Elata | 1.03 ± 0.08 | 0.531 ± 0.010 | 690 ± 70 |
| Pmin | 1.07 ± 0.10 | 0.615 ± 0.013 | 800 ± 20 |
| Pch | 3.33 ± 0.36 | 1.34 ± 0.08 | 1.6 ± 0.1 |
| Elata_Pmin | 1.22 ± 0.107 | 0.877 ± 0.026 | 8.8 ± 0.3 |
| Elata_Pch | 2.25 ± 0.25 | 1.16 ± 0.05 | 1.7 ± 0.2 |
| Pmin_Pch | 4.05 ± 0.20 | 1.62 ± 0.10 | 0.4 ± 0.1 |

Iron reduction was observed in all culture extracts. Elata and Pmin extracts both reduced approximately the same level of iron per mol of phenols (0.531 and 0.615mol iron/mol of phenolic). Pch extracts reduced more iron than the other two fungi per mol of phenolics (1.34mol Fe^2+^/mol of phenolic) reflecting the greater phenolic content (from the F-C results above). Pch extracts also displayed a greater level of reduction per mol of phenolic, reducing more than double the amount of iron compared to the extracts from Elata and Pmin in culture alone. The Elata_Pmin combination showed a significantly increased level of iron reduction when compared to each fungus by itself (p < 0.001, p = 0.001) but the reduction level was not great enough to be considered an additive effect. The combination of Pch_Pmin resulted in more limited total iron reduction when compared to the extracts from each of the fungi when grown separately and added together (p = 0.035, p = 0.005). Iron reduction in the Elata_Pch combined fungal cultures was significantly greater than from Elata alone (p = 0.003), and the reduction level was comparable to the Pch extracts when grown alone (p = 0.06).

### H_2_O_2_ production

H_2_O_2_ is required in the CMF reaction to react with ferrous iron. Although some enzymes are known to generate H_2_O_2_, enzymes are unable to penetrate intact plant cell walls (18, 29, 39-43), and a catalytic source of H_2_O_2_ within the cell wall would promote a more efficient CMF reaction. Redox cycling of chelators with oxygen and transition metals in acidic pH environments is known to generate H_2_O_2_ in solution, thus promoting the production of hydroxyl radicals (28), and we explored whether this mechanism might also be associated with the phenolic components from the three fungi.

Extracts from Elata and Pmin produced the highest levels of H_2_O_2_ (690 and 800mmol H_2_O_2_/mol of phenols) as determined by the FOX assay (Table 1). While Pch and all combinations had detectable levels of H_2_O_2_ generation, they were in the 0.4 to 9mmol range and significantly lower than both Elata and Pmin (Pch vs Elata: p = 0.006, Pch vs Pmin: p < 0.001, Elata vs Pch_Elata: p = 0.006, Pmin vs Pch_Pmin: p = 0.005). Based on the results from the F-C and Ferrozine assay, it was expected that the Elata_Pmin combination extract would show higher levels of H_2_O_2_ production than Elata and Pmin extracts. However, H_2_O_2_ production in extracts from this combination was below 10 mmol, and although still significantly higher than the Pch extract alone (p < 0.001) it was well below the 690 and 800mmol levels seen for the individual cultures of Elata and Pmin respectively. This may be due to the metabolite profile for the combined culture of Elata and Pmin being distinct from the individual cultures and lacking in the primary metabolites seen in the cultures when grown alone (See “Identification of LMW Metabolites” section below).

### Electron Paramagnetic Resonance (EPR)

A DMPO-HO^•^ adduct with its characteristic 4-line spectrum allowed detection and quantification of HO^•^ in solution (Fig. 2). All extracts were found to generate HO^•^ when examined in the EPR studies for radical generation for pH values of both 3.5 and 5.5. The production of HO^•^ was considerably greater at pH 3.5 for most of the treatments tested. It should be noted that only cultures containing Pch reached pH 3-3.5 under the experimental conditions tested. Uniquely, all extracts produced more HO^•^ than a reference catechol compound at the same concentration based on F-C analysis of phenolics. Catechol is typically used as a standard in iron-reduction assays because of its capacity for iron reduction (44).

**FIG 2.**
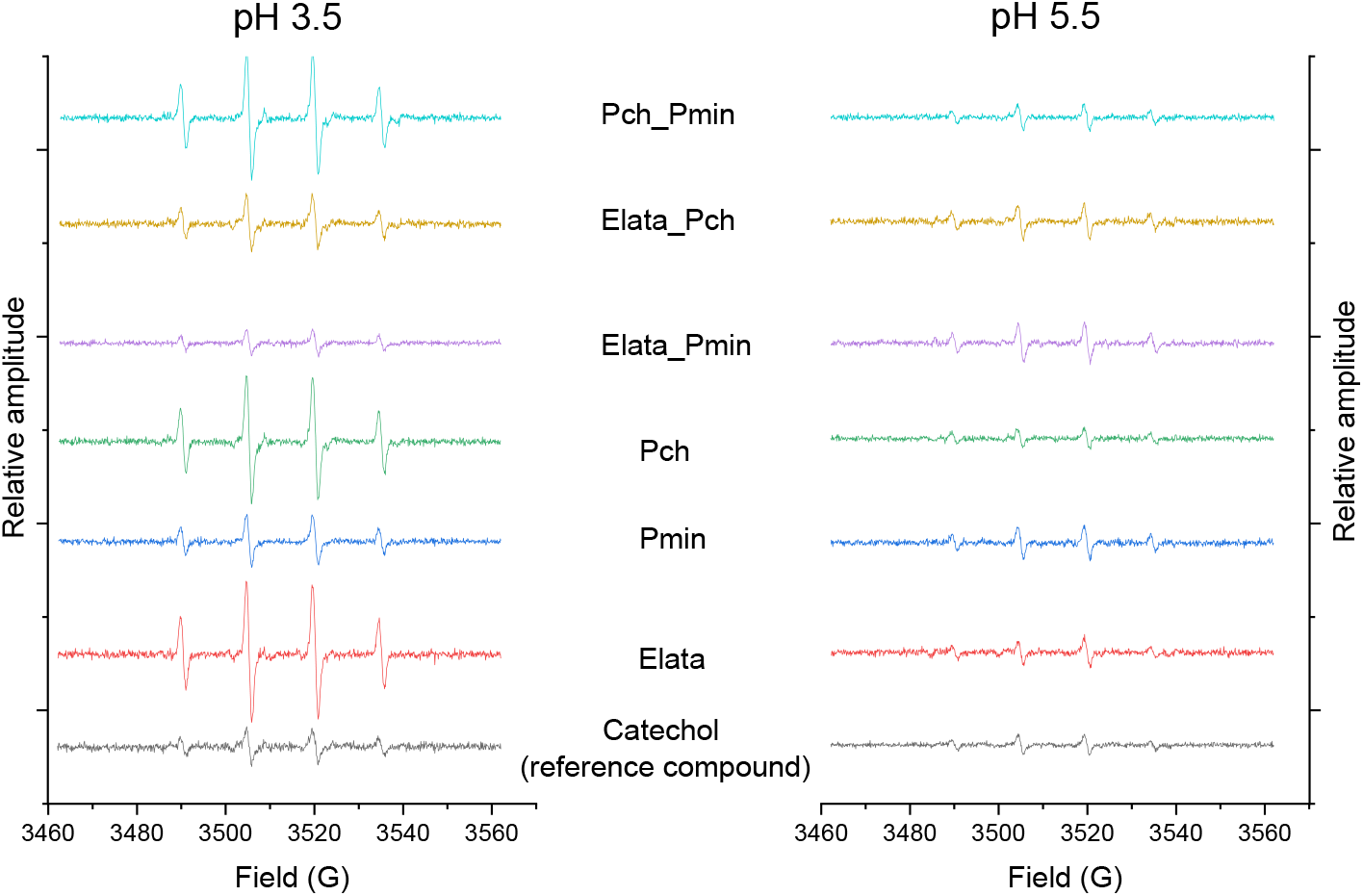
Electron paramagnetic resonance (EPR) spectra of GTD fungal extracts spiked with DMPO to detect hydroxyl radicals. The relative amplitude of each 4-peak spectra reflects the amount of hydroxyl radical produced relative to a catechol standard. Fungi were grown alone, and in consortia, to produce the extracts analyzed in this work.

### Identification of LMW metabolites

Metabolite identification analysis of the three fungi in individual cultures (Fig. S2-S4) showed that several phenolic and non-phenolic compounds were present, with some of them previously reported as having iron reduction capacity (Table 2). Interestingly, all fungi produced small catecholates like 3,4-dihydroxybenzoic acid (Elata), caffeic acid (Pmin) and hydroxycinnamic acids like sinapinic acid (Pch, Pmin) and caffeic acid (Pmin). Catecholates and hydroxycinnamic acids chelate iron and are known for their ability to increase the production of ROS under appropriate conditions (44-46). In addition, Elata also produces pyochelin, a phenolic siderophore and iron reducing compound that is one of the primary siderophores isolated from *Pseudomonas spp*.; and terrein, a non-phenolic iron reducer also found in *Aspergillus terreus* with reported anticarcinogenic activity (47).

**TABLE 2.** Metabolites produced by *E. lata, P. minimum*, and *P. chlamydospora* with previously reported capacity for iron reduction.

| Compound | Formula | Elata | Pmin | Pch | Ref |
| --- | --- | --- | --- | --- | --- |
| Pyochelin | C<sub>14</sub>H<sub>16</sub>N<sub>2</sub>O<sub>3</sub>S<sub>2</sub> | ● |  |  | (48) |
| 3,4-dihydroxybenzoic acid | C<sub>7</sub>H<sub>6</sub>O<sub>4</sub> | ● |  |  | (49) |
| Terrein | C<sub>8</sub>H<sub>10</sub>O<sub>3</sub> | ● |  |  | (50) |
| Sinapic acid methyl ester | C<sub>12</sub>H<sub>14</sub>O<sub>5</sub> |  | ● | ● | (51) |
| Dihydroferulic acid | C<sub>10</sub>H<sub>12</sub>O<sub>4</sub> |  |  | ● | (52) |
| Caffeic acid | C<sub>9</sub>H<sub>8</sub>O<sub>4</sub> |  | ● |  | (53) |
| Gallic acid | C<sub>7</sub>H<sub>6</sub>O<sub>5</sub> |  | ● |  | (54) |
| Homogentisic acid | C<sub>8</sub>H<sub>8</sub>O<sub>4</sub> |  | ● |  | (55) |
| Sinapinic acid | C<sub>11</sub>H<sub>12</sub>O<sub>5</sub> |  | ● |  | (56) |

MS analysis also identified other organic phenols, aldehydes, and carboxylic acids (Table 3) without previously reported iron reduction activity; however, some structures suggest that they could potentially chelate iron. Further experiments with individual model metabolites would be required to demonstrate this, however.

**TABLE 3.**
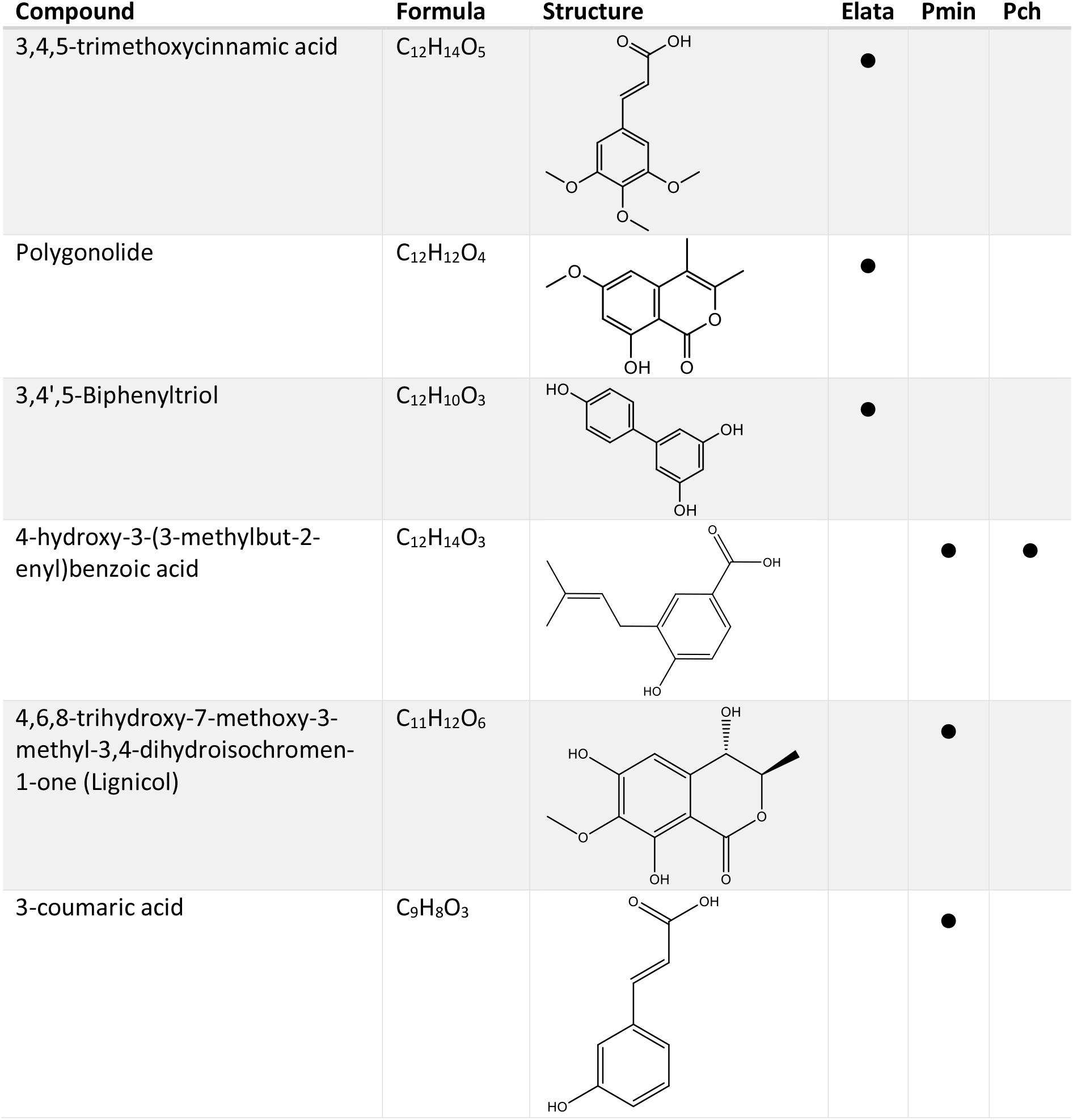
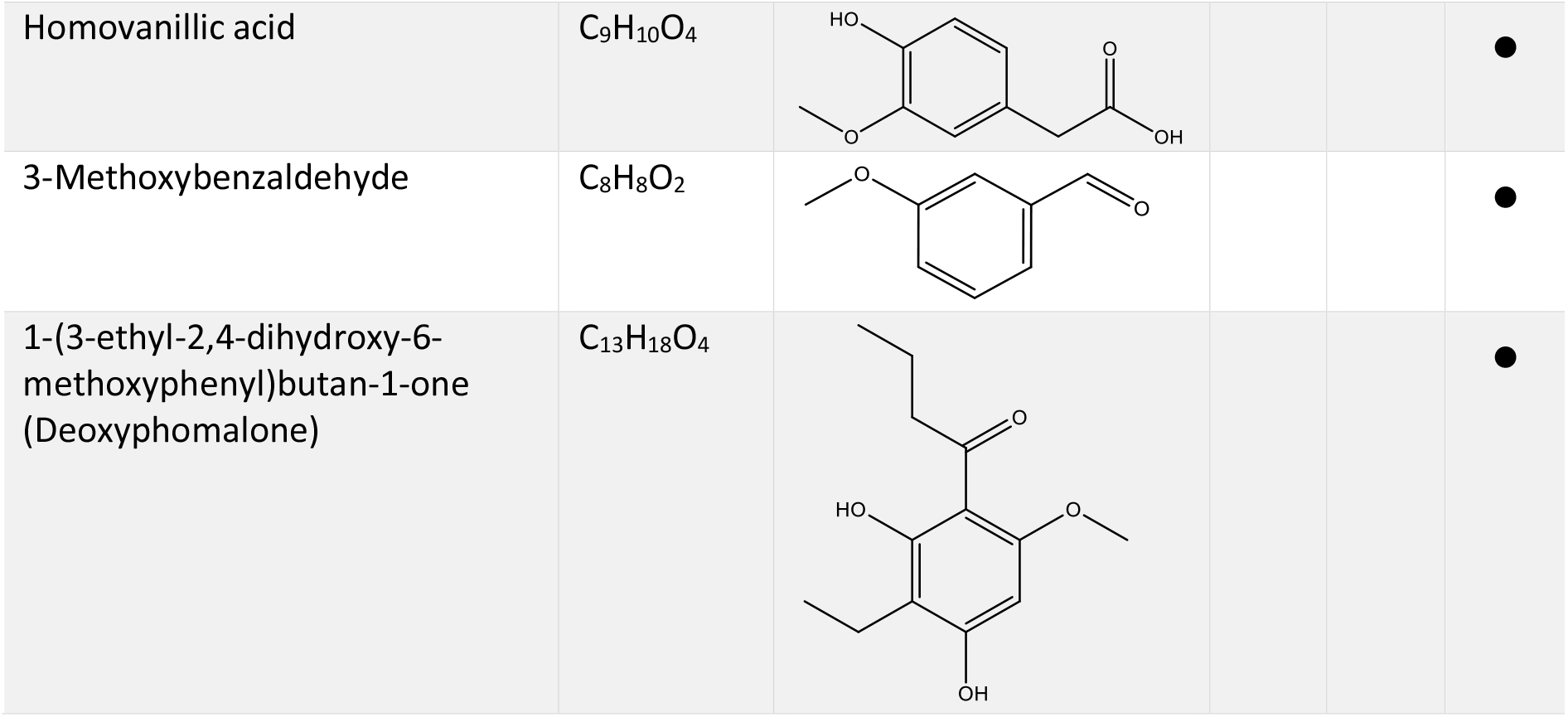
Mass Spectral analysis of phenolic, aldehydes, and carboxylic acid metabolites produced by *E. lata, P. minimum*, and *P. chlamydospore* without reported iron reduction activity.

| Compound | Formula | Structure | Elata | Pmin | Pch |
| --- | --- | --- | --- | --- | --- |
| 3,4,5-trimethoxycinnamic acid | C<sub>12</sub>H<sub>14</sub>O<sub>5</sub> |  | ● |  |  |
| Polygonolide | C<sub>12</sub>H<sub>12</sub>O<sub>4</sub> |  | ● |  |  |
| 3,4',5-Biphenyltriol | C<sub>12</sub>H<sub>10</sub>O<sub>3</sub> |  | ● |  |  |
| 4-hydroxy-3-(3-methylbut-2-enyl)benzoic acid | C<sub>12</sub>H<sub>14</sub>O<sub>3</sub> |  |  | ● | ● |
| 4,6,8-trihydroxy-7-methoxy-3-methyl-3,4-dihydroisochromen-1-one (Lignicol) | C<sub>11</sub>H<sub>12</sub>O<sub>6</sub> |  |  | ● |  |
| 3-coumaric acid | C<sub>9</sub>H<sub>8</sub>O<sub>3</sub> |  |  | ● |  |
| Homovanillic acid | C<sub>9</sub>H<sub>10</sub>O<sub>4</sub> |  |  |  | ● |
| 3-Methoxybenzaldehyde | C<sub>8</sub>H<sub>8</sub>O<sub>2</sub> |  |  |  | ● |
| 1-(3-ethyl-2,4-dihydroxy-6-methoxyphenyl)butan-1-one<br>(Deoxyphomalone) | C<sub>13</sub>H<sub>18</sub>O<sub>4</sub> |  |  |  | ● |

Based on our findings from both the iron reduction and hydrogen peroxide production analyses, we propose that some GTD fungi produce LMW metabolites that are responsible for iron reduction, while other fungi produce LMW metabolites that play a greater role in hydrogen peroxide/oxidant production (Table 1). The LMW extracts from Elata and Pmin possessed relatively limited iron-reduction capacity but produced a significantly greater amount of hydrogen peroxide than Pch. Extracts from the fungal consortia combinations possessed iron reduction capability, but virtually no hydrogen peroxide was observed, including in the combination Elata Pmin. Our MS analysis (Table 2) showed at least five metabolites with iron reduction capacity were identified from Pmin; with Elata and Pch producing 3 and 2 iron reducing metabolites, respectively. We suggest that individual fungi in GTD consortia produce LMW metabolites with specialized and differential functions, with some species taking on greater roles in the production of iron-reducing metabolites while other species produce metabolites with greater capacity for peroxide generation. This may help to explain why GTD pathogenesis is often associated with a consortium of GTD fungi rather than individual fungi.

As detailed in the CMF mechanism for brown rot fungi (19), we proposed that H_2_O_2_ reacts with reduced iron to allow targeted ROS generation such as hydroxyl radicals within plant/wood cell walls to initiate digestion of the lignocellulose components (as part of pathogenesis). In the case of Eutypa dieback and the Esca disease complex, for wood necrosis to occur by a similar mechanism these reactions would also need to occur in tissue that is maintained at a pH ≤ 5.5. This falls into the range of the natural pH of wood cell walls, and our results also show that a pH of ∼5.5 was maintained by individual liquid cultures of Pmin and Elata throughout the growth period of 6 weeks (data not shown). However, in Pch cultures (grown either alone, or in combination with either Pmin or Elata) the pH dropped to 3-3.5. The reduction in pH by Pch may also help explain why different consortia fungi are required in GTD pathogenesis, as pH modification potentially changes the fungal/wood micro-environment to promote the select oxidative chemistries that we report here. Similar oxidative chemistries have also been observed with other catecholate fungal metabolites (19, 25). It has been demonstrated that acidic environments enhance iron reduction by phenolic compounds, and this is observed in our ferrozine assay results (Table 1) where extracts from Pch alone or in consortia reduced more iron than any other fungal extract produced from either individual fungi or consortia growth. The reduction in pH by Pch alone, or in consortia, supports the hypothesis that GTD fungal consortia are necessary for reduction of pH to promote iron reduction as a precursor to the generation of damaging hydroxyl radical generation. This, together with the differential promotion of CMF chemistry by different fungal metabolites, is a novel concept relative to pathogenesis mechanisms associated with consortia fungal activity and grapevine trunk disease that has not previously been reported.

Further exploration of this mechanism could potentially open new alternatives for treatments to prevent or limit yield loss in vineyards. Fig. 3 summarizes our hypothesized mechanism for the initiation of the wood cell wall damage by GTD fungi through HO^•^ generation by LMW metabolite secretion. Lignin depolymerization via this LMW mechanism would promote lignocellulose cell wall damage to produce features of necrosis, and after the initiation of this degradation, would potentially allow extracellular fungal CAZymes to further depolymerize the wood cell wall. In later stages, as vine defenses are overwhelmed, the fungal LMW metabolites would also potentially be transported via the vascular system of the stem to promote further necrosis of cordon wood and ultimately leaf tissue with disease progression.

**FIG 3.**
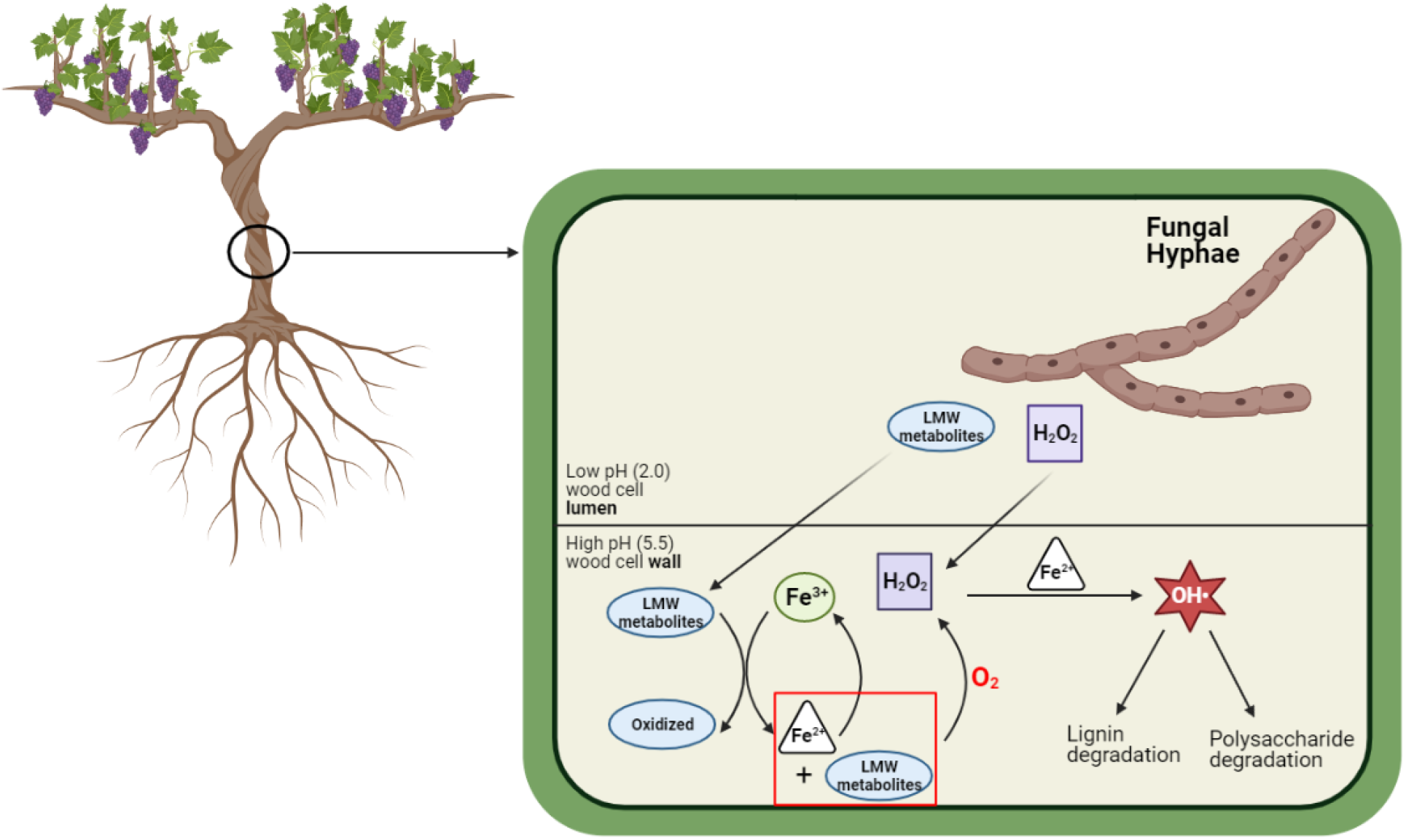
Mechanism for the in-situ generation of Fe^2+^ and H_2_O_2_, and degradation of lignin and cell wall macromolecules by GTD fungi. LMW metabolites and H_2_O_2_ diffuse into the cell wall, where the LMW metabolites sequester Fe^3+^ from the cell wall environment and reduce Fe^3+^ to Fe^2+^. Through the mediated Fenton reaction, Fe^2+^ and H_2_O_2_ react and generate hydroxyl radicals (OH^•^). Images built using Biorender software. Schematic modified from (29).

## CONCLUSIONS

Our data indicates that some GTD fungi preferentially promote iron reduction in culture, while others boost H_2_O_2_ production, suggesting that a mechanism similar to the CMF mechanism employed by brown rot fungi in wood biodegradation may also play a role in GTD pathogenesis. Since both iron reduction and H_2_O_2_ production are required in CMF systems for generation of hydroxyl radicals, this may help to explain why GTDs are often caused by a consortium of fungi rather than individual species. Often these diseases are associated with what has been described previously as non-causative fungi, but the role of LMW in GTD Eutypa and Esca pathogenesis has not been well explored. Our observation of differential function of extracellular LMW metabolites from GTD consortia fungi has not previously been reported, even in brown rot Basidiomycota species where the role of LMW metabolites in wood decay has been established. Thus, this is the first report of the differential action of LMW metabolites promoting the sequential steps of a mediated Fenton chemistry related to pathogenesis in grapevine tissue. We also observed that one fungus in the consortium promoted establishment of a lower pH environment. Pch reduced the pH of the media to 3-4 when grown alone in cultures, but also in consortia with either Elata or Pmin. This pH promotes iron reduction by catecholates, and our observation that Pch reduces pH of the media, while Elata/Pmin maintain pH constant (5.5), may also help to explain why fungal consortia growth is required in many GTDs. Additional research must be conducted to isolate individual LMW metabolites to determine their specific role in GTD fungal consortia ; however, our MS analysis has already identified select metabolites from each of the three fungal species studied which have previously been reported to possess iron-reduction capability. Future research will assess whether these specific LMW metabolites also aid in promoting CMF chemistry and the generation of hydroxyl radicals as a component of pathogenesis.

## ACKNOWLEDGMENTS

The authors would like to thank Prof. Akif Eskalen and Dr. Karina Delfar of the Department of Plant Pathology of the University of California, Davis for providing cultures of *Phaeomoniella chalmydospora, Phaeacremonium minimum* and *Eutypa lata* used in this study. We appreciate the support of Professor Kevin Kittilstved from the Department of Chemistry at the University of Massachusetts, Amherst, in allowing the use of their EPR facilities. Mass Spectral analysis was conducted by Creative biolabs (Shirley, NY).

This research was supported by the National Institute of Food and Agriculture (Multistate Hatch Research Project S1075 1020436-MAS00503), U.S. Department of Agriculture, the Center for Agriculture, Food and the Environment, and the Microbiology Department, University of Massachusetts Amherst. The contents are solely the responsibility of the authors and do not necessarily represent the official views of the USDA or NIFA. The authors also thank the American Vineyard Foundation (project 2019-2275) for their support.

## AUTHOR CONTRIBUTIONS

Conceptualization, Barry Goodell; Methodology, Dana Sebestyen, Gabriel Perez-Gonzalez and Barry Goodell; Validation, Barry Goodell; Formal Analysis, Dana Sebestyen, Gabriel Perez-Gonzalez, Norman Lee; Investigation, Dana Sebestyen and Gabriel Perez-Gonzalez.; Resources, Barry Goodell and Christophe Bertsch.; Writing – Original Draft Preparation, Dana Sebestyen and Gabriel Perez-Gonzalez; Writing – Review & Editing, Dana Sebestyen, Gabriel Perez-Gonzalez, Elsa Petit, Jody Jellison, Laura Mugnai, Christophe Bertsch, Sibylle Farine, Eric Gelhaye, and Barry Goodell; Visualization, Barry Goodell, Dana Sebestyen and Gabriel Perez-Gonzalez; Supervision, Barry Goodell, Gabriel Perez-Gonzalez, Jody Jellison, Elsa Petit; Project Administration, Barry Goodell.; Funding Acquisition, Barry Goodell.

